# AdDeam: A Fast and Scalable Tool for Estimating and Clustering Reference-Level Damage Profiles

**DOI:** 10.1101/2025.03.20.644297

**Authors:** Louis Kraft, Thorfinn Sand Korneliussen, Peter Wad Sackett, Gabriel Renaud

## Abstract

**Motivation:** DNA damage patterns, such as increased frequencies of C*→*T and G*→*A substitutions at fragment ends, are widely used in ancient DNA studies to assess authenticity and detect contamination. In metagenomic studies, fragments can be mapped against multiple references or *de novo* assembled contigs to identify those likely to be ancient. Generating and comparing damage profiles, however, can be both tedious and time-consuming. Although tools exist for estimating damage in single reference genomes and metagenomic datasets, none efficiently cluster damage patterns.

**Results:** To address this methodological gap, we developed `AdDeam`, a tool that combines rapid damage estimation with clustering for streamlined analyses and easy identification of potential contaminants or outliers. Our tool takes aligned aDNA fragments from various samples or contigs as input, computes damage patterns, clusters them and outputs representative damage profiles per cluster, a probability of each sample of pertaining to a cluster as well as a PCA of the damage patterns for each sample for fast visualisation. We evaluated `AdDeam` on both simulated and empirical datasets. `AdDeam` effectively distinguishes different damage levels, such as UDG-treated samples, sample-specific damages from specimens of different time periods, and can also distinguish between contigs containing modern or ancient fragments, providing a clear framework for aDNA authentication and facilitating large-scale analyses.

**Availability and Implementation:** `AdDeam` is publicly available at https://github.com/LouisPwr/AdDeam and can also be installed via Bioconda. It is implemented in Python and C++. All analysis scripts and datasets are available at https://github.com/LouisPwr/AdDeamAnalysis and on Zenodo under: 10.5281/zenodo.15052427.

## 2 Introduction

Validating the authenticity of ancient DNA (aDNA) samples is essential to ensure the reliability of scientific conclusions in evolutionary and archaeological studies [1, 2]. The authentication process relies on the characteristic molecular alterations that occur in DNA molecules. Two primary processes lead to the breakdown and chemical alteration of aDNA: hydrolytic mechanisms cause the degradation of DNA molecules thus resulting in shorter fragments [3, 4], and deamination converts cytosines into uracil molecules. During DNA sequencing, these alterations are interpreted as base substitutions [5]. This phenomenon, known as “damage”, occurs predominantly at the ends of aDNA fragments. In double-stranded aDNA library preparation [6], this leads to elevated frequencies of C*→*T substitutions at the 5^*′*^ end and G*→*A substitutions at the 3^*′*^ end [5, 7]. In contrast, for single-stranded aDNA library preparation [8, 9], only C*→*T substitutions at the 5^*′*^ end are observable. Base substitutions can be partially or nearly completely removed through wet-lab protocols using uracil-DNA glycosylase (UDG) treatment. Full UDG treatment excises uracil residues from aDNA fragments, effectively eliminating damage signatures. While this improves sequence accuracy, it also results in shorter fragment lengths [10]. Partial UDG treatment leaves some uracil residues intact, preserving a limited number of damage sites for authentication [11]. As a result, full UDG treatment produces fragments with no observable damage patterns, whereas partial UDG treatment retains minimal damage signatures. It should be noted that, in certain samples such as those from vertebrates, cytosines in methylated CpG dinucleotides undergo direct deamination to form thymine. Therefore, UDG treatment is ineffective at these sites, resulting in residual C*→*T substitution frequencies even after full treatment [10].

Damage patterns are used to determine whether the DNA being extracted from fossils or ancient environments is indeed ancient in origin [12, 5, 3]. DNA from modern contaminants will not display such damage patterns [13, 14, 15, 16, 17, 18, 19]. In addition, the levels of damage are informative about the conditions in which DNA was preserved, as C*→*T substitutions towards the 5’-ends of aDNA molecules tend to increase over time [20]. When several samples of eukaryotic species like humans are sequenced, researchers routinely screen multiple extractions and determine which carry damage patterns and at which level to determine authenticity [21, 22, 23]. This involves aligning the aDNA fragments to a single reference from the eukaryotic species in question. In the case of metagenomic studies involving *de novo* assembly, damage patterns are used to determine which contigs are likely to be ancient by aligning the aDNA fragments back to the assembly [24, 25, 26, 27, 28]. A contig where the aligned fragments do not display damage patterns is more likely to be a modern contaminant [29]. Furthermore, aDNA fragments from microbes can be aligned to multiple reference genomes and damage patterns are used to determine which species are likely to be present and *bona fine* aDNA [30, 31, 32, 33]. Regardless of which, researchers in aDNA are faced with the task of inspecting large numbers of damage pattern plots to identify outliers.

Several bioinformatic tools generate damage profiles from alignment files, but none are designed to cluster hundreds to thousands of profiles at once to reveal shared damage signatures. Manually inspecting individual damage plots is time-consuming and prone to oversight, and rapid clustering of similar patterns could immediately flag contamination, preservation states, or library-type effects [34, 35]. Moreover, in ancient metagenomic studies involving *de novo* assembly, clustering contig damage profiles can streamline authentication workflows. For example, clusters of contigs that exhibit no damage can serve as calibration sets for statistical tools like `PyDamage` [29]. Visually accessible clusters can inform model parameter selection, such as setting thresholds to distinguish ancient from non-ancient contigs, while clusters with ambiguous signatures flag those contigs for targeted inspection.

Existing tools fall into two broad categories. First, damage profilers such as `mapDamage` [36, 37] and `PMDtools` [38] compute damage profiles from a single reference genome, while `DamageProfiler` [39] can additionally distinguish multiple references within a single file. Second, metagenomic-focused tools and pipelines such as `pyDamage` [29], `metaDamage` [40], `metaDMG` [41], `HOPS` [42], and `aMeta` [43] quantify damage at the contig or taxonomic level, often integrating `mapDamage`, `PMDtools`, and alignment tools like `MALT` [44, 45] into automated workflows.

To our knowledge, only `aRchaic` [34] attempts to cluster fragments by damage patterns, but it is limited to a single predefined reference [46] and is no longer actively maintained. This methodological gap motivated the development of `AdDeam`, which combines (1) efficient computation of damage profiles from one or many BAM files with support for paired-end, single-strand, or mixed libraries and (2) a Gaussian mixture-model clustering framework that groups similar profiles into *k clusters*. *A*dDeam then visualizes both individual and representative damage patterns, enabling rapid, multi-sample screening of damage patterns across diverse references.

Two modes are available: In *classic* mode, `AdDeam` analyses BAM files from different samples aligned to the same reference genome, useful for assessing damage in fragments aligned to a human reference. In *meta* mode, it evaluates fragments from multiple contigs within a single BAM file.

This mode is aimed at researchers who wish to determine which contigs are potentially modern in origin.

Utilizing a Gaussian Mixture Model (GMM), `AdDeam` clusters similar damage profiles, providing both quantitative assessments and clear visualizations. This approach is ideal for organizing aDNA studies with multiple samples or datasets spanning numerous species [23, 31, 30, 47, 48]. `AdDeam` processes large collections of BAM files as input from any reference type (e.g., bacterial, human, or other genomes) as well as BAM files containing alignments to multiple references, such as contigs or grouped reference genomes. It outputs the degree of assignment of the samples or contigs to each cluster and reports the representative damage profile for each cluster.

We demonstrated `AdDeam`’s ability to accurately cluster damage profiles using simulated data and tested its performance on empirical datasets.

## 3 Materials and Methods

In this manuscript, we distinguish between fragments and reads as follows: fragments represent the sequences of degraded aDNA molecules. In paired-end sequencing, each fragment is sequenced from both ends, producing two corresponding reads. Reads are individual observations of the same fragment. However, by trimming adapters and merging paired-end reads, one can reconstruct the most likely sequence representing the original fragment [49]. Consequently, we consistently refer to aDNA fragments when discussing trimmed and merged sequences, such as those used for alignment to a reference.

`AdDeam` consists of two subsequent stages: 1) generating damage profiles from BAM files either for each sample in classic mode or per contig in meta mode 2) clustering these based on their similarity into *k* clusters, where *k* is user-defined. By default, clustering is performed for *k* = 2, 3, and 4. Finally, `AdDeam` outputs a comprehensive report in PDF format as well as tab-separated files for convenient downstream analysis. The workflow of `AdDeam` is illustrated in Figure **1**.

**Figure 1:**
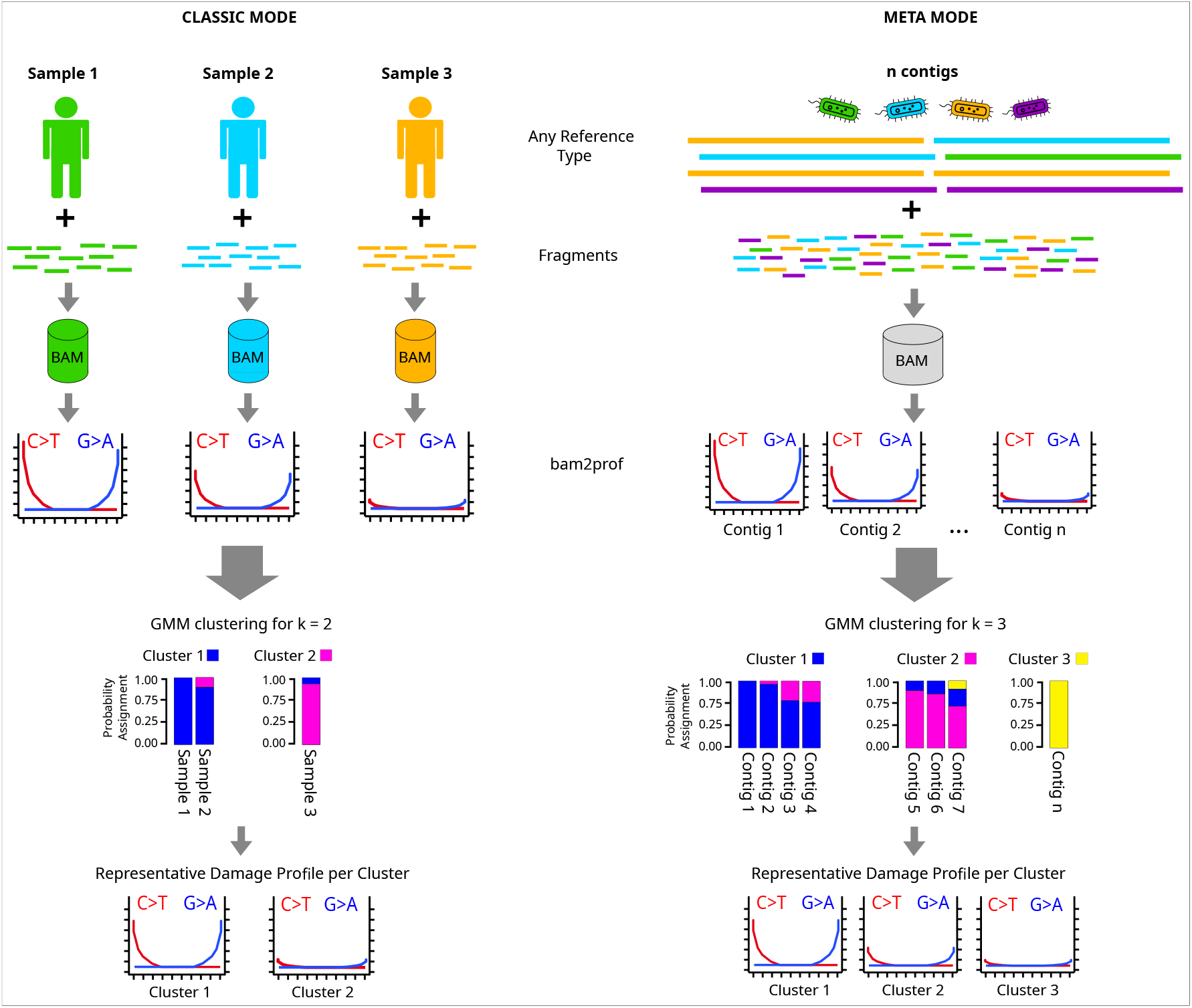
Overview of the two modes of the `AdDeam` workflow: classic mode and meta mode. In classic mode (left), multiple BAM files, each representing different samples mapped to a reference genome (e.g., human), are processed with `bam2prof`, a lightweight C++ program included with `AdDeam`, to generate individual damage profiles. These profiles are then clustered using the GMM algorithm, assigning each sample a probability of belonging to each of the *k* (in this example *k* = 2) clusters. In meta mode (right), one (or more) BAM file containing the original aDNA fragments aligned to a reference containing the assembled contigs is processed. The `bam2prof` program generates damage profiles for each reference within the BAM file. GMM clustering (shown for *k* = 3) then assigns probabilities for each contig’s cluster membership. The bar plots display these stacked membership probabilities by colour. `AdDeam` then outputs the representative damage profile for each cluster.

### 3.1 Stage 1: Damage Profile Generation

We employ `bam2prof`, a tool written in C++, originally developed by Gabriel Renaud and Thorfinn Korneliussen (https://github.com/grenaud/bam2prof), to generate damage profiles from BAM files containing aDNA alignments. `bam2prof` iterates over all alignments and calculates the ratio of observed C*→*T and G*→*A mismatches per position. This stage is independent of the clustering stage and its sole aim is to compute damage patterns. Building upon the original implementation, we have extended the functionality with several key features:

#### Classic mode

Generates a single damage profile per BAM file. This function may be used to analyse a set of BAM files that are aligned to a single reference genome, regardless of whether the reference genome is of bacterial or eukaryotic origin (i.e., human genome with multiple chromosomes), as illustrated in Figure **1**, CLASSIC MODE panel.

#### Meta mode

Produces individual damage profiles for each reference sequence within a BAM file, enabling analysis of damage patterns with each record (be it a contig, scaffold or chromosome) in a reference. This mode is particularly useful for samples where aDNA fragments have been aligned to a database containing multiple reference genomes, such as prokaryotic genomes in metagenomic studies. The result is one damage profile per record in the reference. Additionally, this mode is especially interesting for metagenome assemblies, allowing fragments used for assembly to be aligned back to the resulting contigs, producing a damage profile for each contig (see Figure **1**, META MODE panel).

#### Fast profile generation from large BAM files

BAM files can be very large, which makes computing damage profiles time-intensive. To address this, `bam2prof` implements an iterative convergence check that both accelerates processing and detects unstable (“zig-zag”) profiles. After every *n* aligned fragments, `bam2prof` measures the absolute difference in substitution frequencies (e.g. C*→*T) and terminates early, if this difference falls below a user-defined threshold *d*. Users can adjust *n* and *d* (via `-numAligned` and `-precision`), although the default behaviour ensures processing of all aligned fragments.

This same convergence criterion identifies atypical profiles that are common in ancient metagenomic studies due to low coverage or mismappings. Such non-convergent profiles are written to a separate “notConverged” directory for downstream analysis. A detailed analysis of convergence behaviour (tracking C*→*T changes every 500 aligned fragments) and its impact on damage profile stability is provided in Supplementary Material (Section S3).

#### Python Wrapper for Parallel Processing of Multiple Input Files

To facilitate efficient high-throughput analysis, we developed a Python wrapper around `bam2prof` that enables the parallel processing of multiple BAM files across multiple threads. This wrapper can be invoked with the command:

~~~
addeam-bam2prof.py -i inputBams.txt -o outputDir
~~~

where `inputBams.txt` is a file listing the input BAM files, and `outputDir` specifies the directory for outputting the generated damage profiles. All parameters available for the compiled `bam2prof` program are also supported by the wrapper script and passed on accordingly.

### 3.2 Stage 2: Clustering of Damage Profiles into *k* clusters

The damage profiles generated by `bam2prof` for a single sample are stored as two tab-delimited text files: one representing the substitution rates at the 5’ end of the DNA fragments and the other for the 3’ end. In these files, rows correspond to the first nucleotide positions from each end (5’ or 3’), and columns capture substitution rates for all 12 possible mismatches at each position.

We prepare the profiles for clustering by combining the damage rates. `bam2prof` considers only 5 positions away from the 5’ and 3’ end which get converted into a feature vector of length *p* and therefore, by default, *p* = 10. Specifically, the first five elements of this vector capture the C*→*T substitution frequencies at positions 1 through 5 from the 5’ end, whereas the last five elements represent G*→*A substitution frequencies at positions 5 through 1 from the 3’ end. This approach focuses on substitution patterns that are informative for identifying aDNA, condensing each profile into a single line vector where one line represents a sample (or contig in meta mode). To improve separation during clustering, all values in the vector are transformed using the logarithm of the substitution frequencies. The logarithm scale is useful as a jump from 0% damage to 15% should be more noticeable than a 15% to 30% jump in damage rates. The logarithm space makes smaller values (which are likely to be a contaminant) more salient.

By concatenating the vectors from all profiles, we construct an *m × p* matrix *X*, where each row represents one of *m* samples and each column corresponds to one of the ten informative positions p (five for C→T and five for G*→*A). This matrix serves as input for clustering the damage patterns using a GMM algorithm implemented using `scikit-learn`[50]. Additionally, `AdDeam` provides a `-lib` flag to accommodate different library types in a single run: `-lib paired` uses the C*→*T column from the 5^*′*^ matrix and the G*→*A column from the 3^*′*^ matrix; `-lib single` uses both C*→*T columns (5^*′*^ and 3^*′*^); and `-lib mixed` uses all four columns (C*→*T and G*→*A from both ends) as the input vector for clustering.

Unlike deterministic clustering algorithms such as *k*-means, which assign *ea*ch sample to a single cluster, the GMM algorithm assigns probabilities to each sample of belonging to every cluster, reflecting the model’s uncertainty in classification. This probabilistic approach allows samples to have partial membership across clusters, capturing cases where certain samples do not clearly belong to a single cluster. These ambiguous samples can then be further analysed downstream. However, the number of clusters, *k*, must be specified by the user in advance, as it is an inherent parameter of the GMM algorithm.

The GMM algorithm is a probabilistic model that assumes data is generated from a mixture of *k* Gaussian distributions. Each cluster is represented as a Gaussian distribution with its own mean and covariance structure, which together define the shape and orientation of the cluster. In our application, the number of Gaussian components, *k*, is specified by the user, with default values of *k* = 2, 3, and 4. For each chosen *k*, the GMM estimates the parameters of each Gaussian distribution to best fit the observed data, which in this case are the damage profiles.

The likelihood of observing a sample 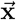 under the GMM is defined as:

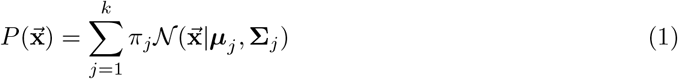

where *π*_*j*_ are the mixing coefficients, ***µ***_*j*_ are the mean vectors, and **Σ**_*j*_ are the covariance matrices of the Gaussian components. Here, 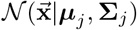 represents the probability density function of the multivariate Gaussian distribution for component *j*, parametrised by its mean vector ***µ***_*j*_ and covariance matrix **Σ**_*j*_. Each Gaussian component represents a potential cluster in the data, with its own probability distribution. Instead of using ***µ***_*j*_ directly as the damage profile, we compute a weighted representative pattern for each cluster by averaging the damage patterns of all samples, weighted by their cluster probabilities. This allows researchers to scan *k* damage profiles rather than hundreds or thousands.

During the development of `AdDeam`, we explored different covariance structures for the GMM algorithm and found that a spherical covariance structure worked best. This structure assumes that each cluster’s covariance matrix is isotropic, meaning that clusters have the same spread in all directions, forming spherical shapes in multi-dimensional space. An advantage of this approach is its computational efficiency, as it requires estimating fewer parameters compared to more flexible structures like a full covariance matrix. The influence of different covariance matrices on clustering outcomes and their applicability to damage patterns is elaborated in the Supplementary Material, Section S2.3. The model is optimized using the Expectation-Maximization algorithm, which iteratively maximizes the likelihood of the observed data under the model, assigning each sample a probability-based cluster membership.

GMM model fitting begins by initializing the component means, which can be chosen at random or set manually. In `AdDeam`, we initialize these means systematically. Here, “values” refers to the minima and maxima of the input matrix *X:*

- For *k* = 2, we set ***µ***_1_ to the minimum values and ***µ***_2_ to the arithmetic mean of minimum and maximum values in linear space.
- For *k* = 3, we add a third mean, ***µ***_3_, set to the maximum values.
- For *k* > 3, additional means are distributed evenly between minimum and maximum values using linear interpolation.

When working with low-coverage metagenomic data, damage profiles can be less reliable if generated from references with minimally aligned fragments [40]. Hence, the Python script that runs the clustering first pre-filters damage profiles based on the number of assigned fragments, ensuring that only reliable profiles are included. Erroneous mappings or low-read evidence can reduce profile accuracy and lead to incorrect clustering outcomes. To address this, profiles generated from references with fewer than a specified threshold of aligned fragments (default: 1,000) are excluded from clustering. Importantly, `bam2prof` also flags non-convergent (“zig-zag”) profiles and writes them to a separate “notConverged” directory. These unstable profiles are not passed to the clustering algorithm unless the user explicitly points the script at their directory.

### 3.3 Output and Visualisation

After clustering the damage profiles, each sample is assigned probabilities indicating its membership in each of the *k* clusters. However, some samples may exhibit nearly uniform probability distributions across multiple clusters, making their classification ambiguous.

To aid interpretation and identify potential outliers, we include a visualization step that projects the clustered damage profiles into a lower-dimensional space using Principal Component Analysis (PCA). Analogous to clustering in logarithmic space, we also perform the PCA on the logarithm of the substitution frequencies. The two most significant principal components are plotted, with each sample colour-coded according to its highest GMM cluster assignment. This provides an intuitive way to assess cluster separation and detect samples with uncertain assignments.

Additionally, we generate probability plots illustrating each sample’s cluster membership probabilities and compute representative damage profiles for each identified cluster as weighted averages based on GMM assignment probabilities.

For downstream analysis, all results, including sample identifiers, mapped read counts, cluster assignment probabilities, and Euclidean distances to GMM cluster centers in PCA space, are exported as tab-separated files.

## 4 Results

We focus on this simulation to demonstrate the feasibility of our clustering approach. As there are no existing tools that appear to cluster damage parameters in such a way, direct benchmarking is difficult. Most available methods either generate damage profiles or classify the probability that, for example, a contig is ancient [36, 37, 38, 39, 40, 43, 42, 41, 29], but they do not cluster damage profiles, which is one of the core functionalities of `AdDeam`. While `aRchaic` [34] could also perform clustering, it does so under the assumption of a single predefined reference, without distinguishing between multiple references within a BAM file. As a result, it cannot capture damage variation across multiple references, which is crucial in metagenomic studies. Additionally, `aRchaic` is no longer maintained, and despite extensive efforts, we were unable to install it successfully. We nevertheless include in the Supplementary Material (Section S2) two comparative analyses: one benchmarking `AdDeam` against the original `aRchaic` results from Al-Asadi *et al*. [34], and another comparing `AdDeam` clusters to `PyDamage` [29] classifications.

### 4.1 Clustering of Simulated Damage Profiles

The damage patterns used for simulating ancient reads served as the ground truth, exhibiting distinct substitution rate intervals for no damage, mid-damage, and high damage (see details in Supplementary Material). However, the estimated damage profiles generated by `bam2prof` inevitably deviate from this ground truth due to alignment biases. These biases arise because not all fragments are perfectly mapped to the reference, leading to overlapping damage rate intervals after estimation. This effect is further detailed in Supplementary Material, Section “Simulated Datasets: Additional Information”, and Figure S1.

In order to evaluate `AdDeam`’s ability to correctly cluster similar damage patterns, we used simulated datasets in which the ground truth damage levels were known to us but not to the algorithm. After running `AdDeam`’s clustering module, we compared its output with the ground truth. From a simulated dataset of 890 samples, we selected a subset comprising 290 samples with no damage, 145 samples with mid damage, and 29 samples with high damage to reflect the empirical heterogeneity of damage patterns. In total, 464 samples were analysed.

Figure **2**, upper panel “Ground Truth of Simulation” illustrates the distribution of samples across the three damage levels that were simulated and subsequently used as input for `AdDeam`.

On the left side of Figure **2**, “Output 1: Gaussian Mixture Model Cluster Assignment,” the automatically generated clustering results for *k* = 2 and *k* = 3 are *show*n.

**Figure 2:**
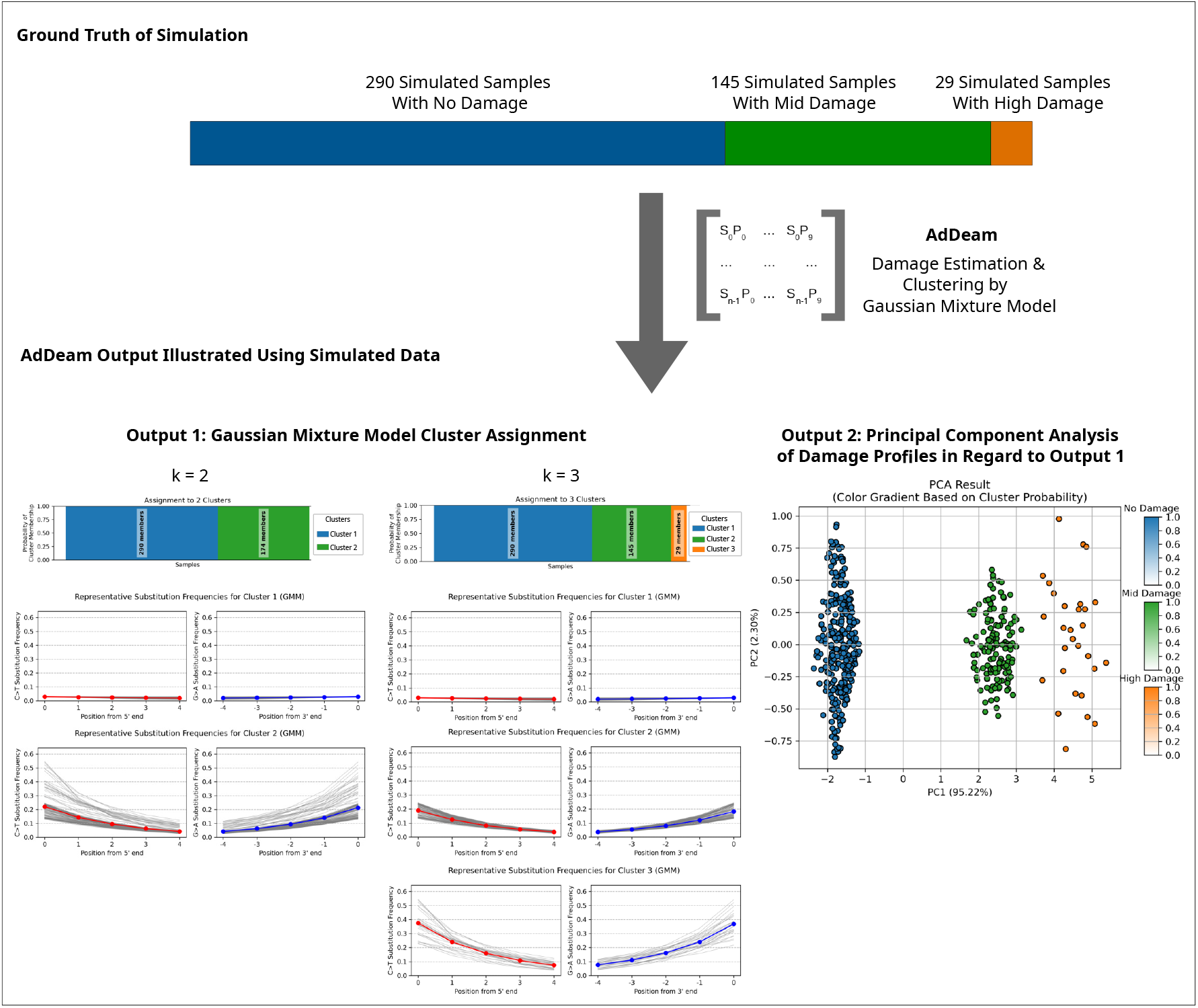
Clustering of simulated damage profiles. Upper panel: Ground truth distribution of samples across the three simulated damage levels used as input for `AdDeam`. The damage plots above the bars illustrate the range of substitution rates that individual samples within each damage level carry. Lower panel left: GMM algorithm clustering results for *k* = 2 and *k* = 3. For *k* = 2, no-damage profiles form one cluster, while mid- and high-damage profiles are combined into a second cluster. For *k* = 3, all profiles are correctly assigned to their respective categories. Lower panel right: PCA of damage profiles for *k* = 3. The plot reveals three distinct clusters, with colour-coding derived from the GMM algorithm.

For *k* = 2, `AdDeam` grouped all no-damage profiles into a single cluster, while mid- and high-damage profiles were combined into a second cluster. Representative damage profiles for each cluster are also provided. The representative profile for the no-damage cluster closely aligns with the expected near-zero substitution frequencies. The representative profile for the combined mid- and high-damage cluster is more similar to the mid-damage profile. This results from the cluster being dominated by 145 mid-damage profiles, compared to only 29 high-damage profiles, making the representative profile weighted accordingly.

For *k* = 3, `AdDeam` successfully assigned all profiles to their correct categories. The representative profiles for the three clusters were distinct and aligned with the expected patterns of no-damage, mid-damage, and high-damage.

On the right side of Figure **2**, “Output 2: Principal Component Analysis of Damage Profiles in Regard to Output 1” presents a PCA plot of the clustering results for *k* = 3. The plot reveals three distinct clusters, visually apparent from the data point distributions and supported by colour coding from the GMM algorithm. No-damage, mid-damage, and high-damage profiles each form separate clusters. The uniform colour within each group indicates perfect consistency in the classification since each cluster only contains samples of the same simulated ground truth of damage. This colour-coded PCA plot is automatically generated by `AdDeam` and provided to the user.

### 4.2 Empirical Analysis: Mapping Ancient Metagenomic Fragments to Core Oral Microbiome Genera

In order to evaluate `AdDeam`’s ability to cluster damage patterns in empirical datasets, we analysed fragments from 49 ancient metagenome datasets originally presented by Fellows Yates *et al*.[30]. These datasets were selected because key characteristics such as sample age and UDG treatment were known, providing metrics to validate `AdDeam`’s clustering performance. The samples, which were extracted from dental calculus, were obtained from various species, including howler monkeys, gorillas, chimpanzees, Neanderthals, ancient modern humans, and present-day modern humans.

We mapped the 49 samples against the reference genomes of four bacterial species: *Tannerella forsythia, Porphyromonas gingivalis, Treponema denticola*, and *Fretibacterium fastidiosum*. These species were chosen because they were described as “core genera” in the study by Fellows Yates *et al*.[30], which defines a core genus as one found in at least two-thirds of the populations within a given host genus. These bacteria are highly prevalent across all host genera in the source study, including *Alouatta* (Howler monkey), *Gorilla, Pan troglodytes* (Chimpanzee), and *Homo Sapiens*. In modern humans, they are strongly associated with periodontal disease [30], and analysing their ancient genomes provides valuable insights into the evolution of the oral microbiome and its relationship to host health.

Therefore, the damage profiles for these four core genera species are of particular interest: their presence can be confidently inferred if ancient damage patterns are observed, and the effectiveness of UDG treatment can be assessed, ensuring the removal of base substitutions crucial for downstream analyses.

The BAM files containing the alignments of the 49 samples were analysed using `AdDeam` in “classic mode”, which means that a single damage profile was reported for the four “core genera”. Figure **3** shows the clustering results for *k* = 4 and the metadata associated with the samples, including cluster assignment, sample source, sample age, and UDG treatment. We saw that *k* = 4 provided a clear separation of the different samples. The left panel displays the probability assignment of each sample to a cluster. The uniform colour of each bar indicates that all samples were assigned to a cluster with probabilities close to 100%. The right panel shows the representative damage patterns for each cluster.

**Figure 3:**
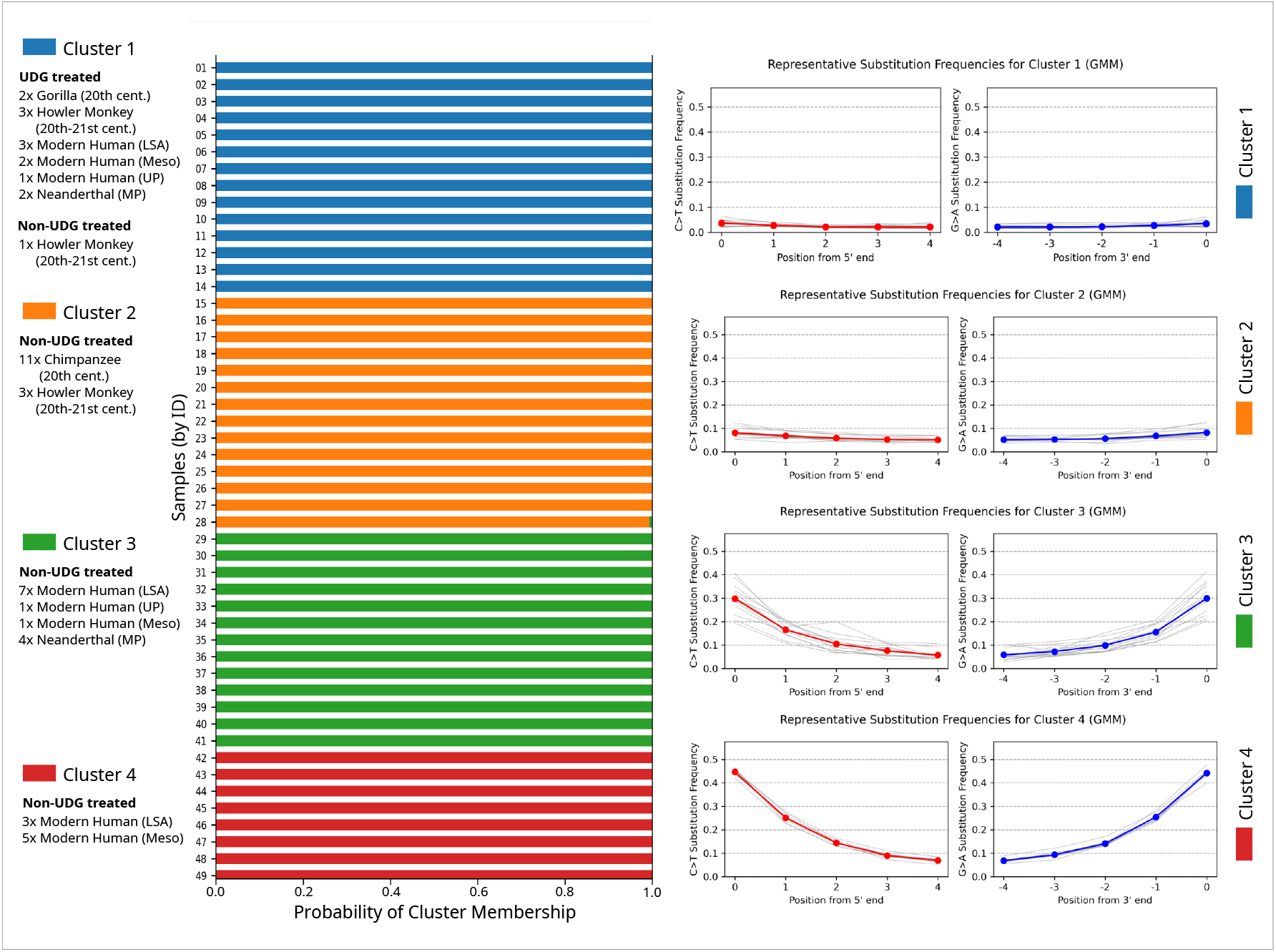
Clustering results for 49 samples aligned against the genomes of four bacterial species that represent the “core genera” of oral microbiomes. A single damage profile was obtained per sample using the alignments to these four genomes which are usually present in oral microbiome. Left: Probability assignments to each cluster (*k* = 4). Right: Representative damage profiles for each cluster. `AdDeam` was run in “classic mode”, generating a single damage profile for all four bacterial species.

Clusters 1 and 2 exhibit representative damage profiles close to zero (cluster 1) and with minimal damage (maximum 5%) for cluster 2. In contrast, the representative profiles for clusters 3 and 4 display more pronounced damage patterns. Cluster 3 shows damage around 30% at the first position, while cluster 4 shows high damage with over 40% at the first position.

To provide additional context, Figure **3** also includes metadata of the samples such as the sample source, the period they originate from and whether they were UDG-treated or not. A Table that also includes the samples codes from Fellows-Yates *et al*. [30] can be found in the Supplementary Material.

Figure **3** reveals that cluster 1 primarily contains all UDG-treated samples from all time periods, with a single exception. This exception is a sample from a howler monkey (*Alouatta*) dated to the 20th-21st century. The damage profile of this outlier shows exceptionally low substitution frequencies, consistent with the UDG-treated samples. This explains its assignment to cluster 1. The output files of `AdDeam` including the damage profiles of all samples are accessible at https://github.com/LouisPwr/AdDeamAnalysis.

Cluster 2 consists exclusively of non-UDG-treated samples from relatively modern time periods (20th-21st century). In contrast, clusters 3 and 4 only include samples from older periods, specifically the Mesolithic, Upper Paleolithic, Middle Paleolithic, and Later Stone Age.

### 4.3 Empirical Analysis: Clustering Damage Profiles from Assembled Contigs

As described in the Introduction `AdDeam` can generate damage profiles for individual records (i.e contigs, scaffolds or chromosomes) in a reference genome using `bam2prof`. This feature enables an interesting use case: clustering damage profiles derived from multiple reference sequences, such as those obtained through *de novo* assembly of ancient metagenomes. The fragments can be realigned to the contigs and determine which are potential contaminants.

To explore this, we analysed a co-assembly strategy by combining one modern sample (Sample ID VLC) and two ancient samples (Sample IDs TAF and EMN) all from Fellows Yates *et al*. [30]. The combined dataset was assembled using `MEGAHIT`[51] and the resulting contigs were filtered to have lengths of at least 10,000 bp. The original fragments which were used for assembly were then mapped to these contigs, and the alignment files were analysed using `AdDeam`. The ENA accession numbers included in the FASTQ headers of each sample enabled us to trace which fragments originated from each sample. Therefore, we could determine for each cluster the fraction of sample-specific fragments that aligned to the contigs. This helped us inferring the composition of the contigs, showing whether they were mostly assembled from modern or ancient fragments.

The results of this analysis are shown in Figure **4**. The left panel displays a PCA plot, where contigs are clustered into three groups (*k* = 3) based on their damage profiles. To validate whether the contigs clustered together match the composition of the reads from which they were assembled, we counted the fragments aligning to contigs in each cluster and calculated the fraction coming from each sample (VLC for modern, TAF for Later Stone Age, and EMN for Upper Paleolithic). The right panel of Figure **4** shows that contigs in cluster 1 have mapped almost exclusively modern fragments (*≥* 99%), contigs in cluster 3 have mapped almost exclusively ancient fragments, and contigs in cluster 2 represent a mix of both. These results align with the representative damage patterns shown.

**Figure 4:**
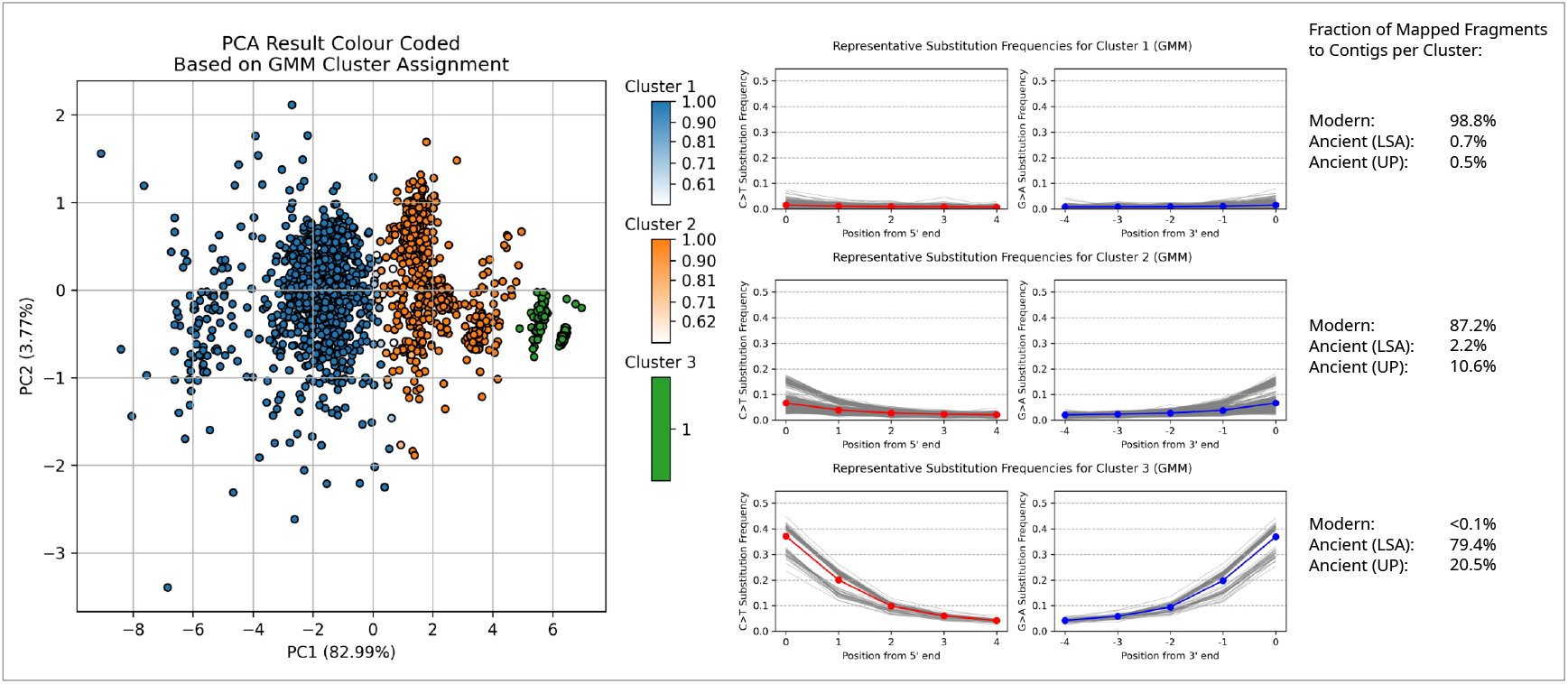
Classifying contigs based on their damage profiles from co-assembled fragments of modern and ancient sources. Three samples from Fellows Yates *et al*. [30] were merged and assembled using `MEGAHIT`[51]. Fragments were mapped to contigs that are over 10,000 bp in length, followed by analysis with `AdDeam` to generate damage reports. The results demonstrate the ability to differentiate between modern and ancient contigs. Clustering analysis (*k*=3) was applied to the VLC (21st century), TAF (Later Stone Age), and EMN (Upper Palaeolithic) datasets. The PCA plot is based on damage profiles from contigs over 10,000 bp, with clusters coloured according to the clustering results. The legend next to each cluster shows the representative damage profile (in order) and the proportion of sample-specific fragments aligning to the contigs in each cluster (Modern: VLC, Ancient (LSA): TAF, Ancient (UP): TAF).

The PCA plot in Figure **4** also shows that some contigs between cluster 1 and 2 are not clearly assigned to a single cluster by the GMM. These contigs are represented by lighter gradient colours, representing lower probabilities of cluster membership. This uncertainty in cluster assignment is consistent with their intermediate positions in the PCA plot, where these data points visually appear between cluster 1 and 2.

Execution of both `AdDeam` commands completed in under 2.5 minutes and required less than 400 MB of memory, as detailed in the Supplementary Material.

## 5 Discussion

We developed `AdDeam`, a versatile and user-friendly tool for estimating DNA damage and clustering damage profiles. Designed to streamline the analysis and comparison of damage patterns, `AdDeam` fills a methodological gap when working with large datasets, such as those containing numerous samples or alignment files with multiple references. As publicly accessible aDNA datasets continue to grow, including million-year-old specimens with highly variable damage patterns [23, 52], the need for efficient and versatile screening tools becomes ever more critical.

Because the GMM requires the user to specify the number of clusters, *k*, it is important to consider what each choice represents. Intuitively, *k* = 2 aligns with the fundamental question in aDNA research of damaged versus undamaged DNA molecules [53]. However, we have shown that values of k > 2 offer benefits as well. For example, *k* = 3 streamlines quality control by partitioning modern (negative control), non-UDG, and UDG-treated libraries (Supplementary Material, Section S2). *De novo* assembled contigs in ancient metagenomic studies can exhibit a broad spectrum of damage. In such settings, intermediate clusters (e.g., *k* = 3 in Figure **4**) can capture contigs with ambiguous profiles, flagging them for further investigation. Therefore, clustering with *k* > 2 can guide authentication workflows, for example, by using low-damage clusters as a visual baseline for adjusting significance thresholds of tools like `PyDamage` [29] (see Supplementary Material Section S2.2).

Future work on clustering damage profiles should further explore algorithms that account for positional dependencies. Our default spherical-covariance GMM assumes independence between read-position features, neglecting true sequential dependencies. However, in our benchmarks, GMMs with full covariance matrices performed poorly whereas DTW-based (Dynamic Time Warping) clustering performed similar to the spherical GMM but still failed to outperform it consistently (Supplementary Material, Section S2.3). Investigating further methods to integrate positional dependencies could enhance clustering of complex datasets (e.g., mapped fragments to thousands of *de novo* assembled contigs) by revealing additional patterns and enabling more fine-grained clusters.

During development of `AdDeam`, we considered whether clustering damage profiles might reveal preservation or depositional “fingerprints” beyond simple authentication. While Sawyer *et al*. [20] noted that variable and often unknown preservation conditions make it unlikely for DNA damage to correlate directly with sample age, no large-scale study has yet tested whether damage-profile signatures are associated with geographic or environmental factors. By enabling rapid clustering, visualization, and versatile application to diverse datasets, `AdDeam` lays the groundwork for future explorations of the ecological and spatial dimensions of aDNA preservation.

## 6 Declarations

### Author Contributions

G.R. conceived the project. L.K. and G.R. designed and developed the method. G.R. and T.S.K. developed the first C++ version of `bam2prof`. L.K. further modified `bam2prof`. L.K. implemented the method and tested it. L.K. conducted all tests.

## References

[1] R. E. Green, A. W. Briggs, J. Krause, K. Prüfer, H. A. Burbano, M. Siebauer, M. Lachmann, and S. Pääbo, “The Neandertal genome and ancient DNA authenticity,” The EMBO Journal, vol. 28, p. 2494–2502, Aug. 2009.

[2] C. Der Sarkissian, I. M. Velsko, A. K. Fotakis, Å. J. Vågene, A. Hübner, and J. A. Fellows Yates, “Ancient Metagenomic Studies: Considerations for the Wider Scientific Community,” mSystems, vol. 6, 12 2021.

[3] M. Hofreiter, D. Serre, H. N. Poinar, M. Kuch, and S. Pääbo, “Ancient DNA,” Nature Reviews Genetics, vol. 2, p. 353–359, May 2001.

[4] S. Pääbo, “Ancient DNA: extraction, characterization, molecular cloning, and enzymatic amplification.,” Proceedings of the National Academy of Sciences, vol. 86, p. 1939–1943, Mar. 1989.

[5] A. W. Briggs, U. Stenzel, P. L. F. Johnson, R. E. Green, J. Kelso, K. Prüfer, M. Meyer, J. Krause, M. T. Ronan, M. Lachmann, and S. Pääbo, “Patterns of damage in genomic DNA sequences from a Neandertal,” Proceedings of the National Academy of Sciences, vol. 104, p. 14616–14621, 9 2007.

[6] A. W. Briggs and P. Heyn, Preparation of Next-Generation Sequencing Libraries from Damaged DNA, p. 143–154. Humana Press, Dec. 2011.

[7] J. Dabney, M. Meyer, and S. Paabo, “Ancient DNA Damage,” Cold Spring Harbor Perspectives in Biology, vol. 5, p. a012567–a012567, 5 2013.

[8] M.-T. Gansauge, T. Gerber, I. Glocke, P. Korlević, L. Lippik, S. Nagel, L. M. Riehl, A. Schmidt, and M. Meyer, “Single-stranded DNA library preparation from highly degraded DNA usingT4DNA ligase,” Nucleic Acids Research, p. gkx033, Jan. 2017.

[9] J. D. Kapp, R. E. Green, and B. Shapiro, “A Fast and Efficient Single-stranded Genomic Library Preparation Method Optimized for Ancient DNA,” Journal of Heredity, vol. 112, p. 241–249, Mar. 2021.

[10] A. W. Briggs, U. Stenzel, M. Meyer, J. Krause, M. Kircher, and S. Pääbo, “Removal of deaminated cytosines and detection of in vivo methylation in ancient DNA,” Nucleic Acids Research, vol. 38, p. e87–e87, Dec. 2009.

[11] N. Rohland, E. Harney, S. Mallick, S. Nordenfelt, and D. Reich, “Partial uracil–DNA–glycosylase treatment for screening of ancient DNA,” Philosophical Transactions of the Royal Society B: Biological Sciences, vol. 370, p. 20130624, Jan. 2015.

[12] M. Hofreiter, V. Jaenicke, D. Serre, A. von Haeseler, and S. Pääbo, “DNA sequences from multiple amplifications reveal artifacts induced by cytosine deamination in ancient DNA,” Nucleic Acids Research, vol. 29, pp. 4793–4799, Dec. 2001.

[13] M. Richards, B. Sykes, and R. Hedges, “Authenticating DNA Extracted From Ancient Skeletal Remains,” Journal of Archaeological Science, vol. 22, p. 291–299, Mar. 1995.

[14] M. L. Sampietro, M. T. P. Gilbert, O. Lao, D. Caramelli, M. Lari, J. Bertranpetit, and C. Lalueza-Fox, “Tracking down Human Contamination in Ancient Human Teeth,” Molecular Biology and Evolution, vol. 23, p. 1801–1807, June 2006.

[15] J. Krause, A. W. Briggs, M. Kircher, T. Maricic, N. Zwyns, A. Derevianko, and S. Pääbo, “A Complete mtDNA Genome of an Early Modern Human from Kostenki, Russia,” Current Biology, vol. 20, p. 231–236, Feb. 2010.

[16] O. Handt, M. Krings, R. H. Ward, and S. Pääbo, “The retrieval of ancient human DNA sequences,” American Journal of Human Genetics, vol. 59, pp. 368–376, Aug. 1996.

[17] C. Warinner, A. Herbig, A. Mann, J. A. Fellows Yates, C. L. Weiß, H. A. Burbano, L. Orlando, and J. Krause, “A robust framework for microbial archaeology,” Annual Review of Genomics and Human Genetics, vol. 18, pp. 321–356, Aug. 2017.

[18] F. M. Key, C. Posth, J. Krause, A. Herbig, and K. I. Bos, “Mining metagenomic data sets for ancient DNA: Recommended protocols for authentication,” Trends in Genetics, vol. 33, pp. 508–520, Aug. 2017.

[19] G. Renaud, V. Slon, A. T. Duggan, and J. Kelso, “Schmutzi: estimation of contamination and endogenous mitochondrial consensus calling for ancient DNA,” Genome Biology, vol. 16, Oct. 2015.

[20] S. Sawyer, J. Krause, K. Guschanski, V. Savolainen, and S. Pääbo, “Temporal Patterns of Nucleotide Misincorporations and DNA Fragmentation in Ancient DNA,” PLOS ONE, vol. 7, p. e34131, Mar. 2012.

[21] D. Pandey, M. Harris, N. R. Garud, and V. M. Narasimhan, “Leveraging ancient DNA to uncover signals of natural selection in Europe lost due to admixture or drift,” Nature Communications, vol. 15, Nov. 2024.

[22] M. E. Allentoft, M. Sikora, A. Fischer, K.-G. Sjögren, A. Ingason, R. Macleod, A. Rosengren, B. Schulz Paulsson, M. L. S. Jørkov, M. Novosolov, J. Stenderup, T. D. Price, M. Fischer Mortensen, A. B. Nielsen, M. Ulfeldt Hede, L. Sørensen, P. O. Nielsen, P. Rasmussen, T. Z. T. Jensen, A. Refoyo-Martínez, E. K. Irving-Pease, W. Barrie, A. Pearson, B. Sousa da Mota, F. Demeter, R. A. Henriksen, T. Vimala, H. McColl, A. Vaughn, L. Vinner, G. Renaud, A. Stern, N. N. Johannsen, A. D. Ramsøe, A. J. Schork, A. Ruter, A. B. Gotfredsen, B. Henning Nielsen, E. Brinch Petersen, E. Kannegaard, J. Hansen, K. Buck Pedersen, L. Pedersen, L. Klassen, M. Meldgaard, M. Johansen, O. C. Uldum, P. Lotz, P. Lysdahl, P. Bangsgaard, P. V. Petersen, R. Maring, R. Iversen, S. Wåhlin, S. Anker Sørensen, S. H. Andersen, T. Jørgensen, N. Lynnerup, D. J. Lawson, S. Rasmussen, T. S. Korneliussen, K. H. Kjær, R. Durbin, R. Nielsen, O. Delaneau, T. Werge, K. Kristiansen, and E. Willerslev, “100 ancient genomes show repeated population turnovers in Neolithic Denmark,” Nature, vol. 625, p. 329–337, Jan. 2024.

[23] K. H. Kjær, M. Winther Pedersen, B. De Sanctis, B. De Cahsan, T. S. Korneliussen, C. S. Michelsen, K. K. Sand, S. Jelavić, A. H. Ruter, A. M. A. Schmidt, K. K. Kjeldsen, A. S. Tesakov, I. Snowball, J. C. Gosse, I. G. Alsos, Y. Wang, C. Dockter, M. Rasmussen, M. E. Jørgensen, B. Skadhauge, A. Prohaska, J. Å. Kristensen, M. Bjerager, M. E. Allentoft, E. Coissac, A. Rouillard, A. Simakova, A. Fernandez-Guerra, C. Bowler, M. Macias-Fauria, L. Vinner, J. J. Welch, A. J. Hidy, M. Sikora, M. J. Collins, R. Durbin, N. K. Larsen, and E. Willerslev, “A 2-million-year-old ecosystem in Greenland uncovered by environmental DNA,” Nature, vol. 612, pp. 283–291, 12 2022.

[24] H. P. Hodgins, P. Chen, B. Lobb, X. Wei, B. J. M. Tremblay, M. J. Mansfield, V. C. Y. Lee, P.-G. Lee, J. Coffin, A. T. Duggan, A. E. Dolphin, G. Renaud, M. Dong, and A. C. Doxey, “Ancient Clostridium DNA and variants of tetanus neurotoxins associated with human archaeological remains,” Nature Communications, vol. 14, Sept. 2023.

[25] P. Rozwalak, J. Barylski, Y. Wijesekara, B. E. Dutilh, and A. Zielezinski, “Ultracon-served bacteriophage genome sequence identified in 1300-year-old human palaeofaeces,” Nature Communications, vol. 15, 1 2024.

[26] M. Klapper, A. Hübner, A. Ibrahim, I. Wasmuth, M. Borry, V. G. Haensch, S. Zhang, W. K. Al-Jammal, H. Suma, J. A. Fellows Yates, J. Frangenberg, I. M. Velsko, S. Chowdhury, R. Herbst, E. V. Bratovanov, H.-M. Dahse, T. Horch, C. Hertweck, M. R. González Morales, L. G. Straus, I. Vilotijevic, C. Warinner, and P. Stallforth, “Natural products from reconstructed bacterial genomes of the Middle and Upper Paleolithic,” Science, vol. 380, p. 619–624, 5 2023.

[27] J. C. Brealey, H. G. Leitão, T. van der Valk, W. Xu, K. Bougiouri, L. Dalén, and K. Guschanski, “Dental Calculus as a Tool to Study the Evolution of the Mammalian Oral Microbiome,” Molecular Biology and Evolution, vol. 37, p. 3003–3022, 5 2020.

[28] M. C. Wibowo, Z. Yang, M. Borry, A. Hübner, K. D. Huang, B. T. Tierney, S. Zimmerman, F. Barajas-Olmos, C. Contreras-Cubas, H. García-Ortiz, A. Martínez-Hernández, J. M. Luber, P. Kirstahler, T. Blohm, F. E. Smiley, R. Arnold, S. A. Ballal, S. J. Pamp, J. Russ, F. Maixner, O. Rota-Stabelli, N. Segata, K. Reinhard, L. Orozco, C. Warinner, M. Snow, S. LeBlanc, and A. D. Kostic, “Reconstruction of ancient microbial genomes from the human gut,” Nature, vol. 594, p. 234–239, 5 2021.

[29] M. Borry, A. Hübner, A. B. Rohrlach, and C. Warinner, “PyDamage: automated ancient damage identification and estimation for contigs in ancient DNA de novo assembly,” PeerJ, vol. 9, p. e11845, July 2021.

[30] J. A. Fellows Yates, I. M. Velsko, F. Aron, C. Posth, C. A. Hofman, R. M. Austin, C. E. Parker, A. E. Mann, K. Nägele, K. W. Arthur, J. W. Arthur, C. C. Bauer, I. Crevecoeur, C. Cupillard, M. C. Curtis, L. Dalén, M. Díaz-Zorita Bonilla, J. C. Díez Fernández-Lomana, D. G. Drucker, E. Escribano Escrivá, M. Francken, V. E. Gibbon, M. R. González Morales, A. Grande Mateu, K. Harvati, A. G. Henry, L. Humphrey, M. Menéndez, D. Mihailović, M. Peresani, S. Rodríguez Moroder, M. Roksandic, H. Rougier, S. Sázelová, J. T. Stock, L. G. Straus, J. Svoboda, B. Teßmann, M. J. Walker, R. C. Power, C. M. Lewis, K. Sankaranarayanan, K. Guschanski, R. W. Wrangham, F. E. Dewhirst, D. C. Salazar-García, J. Krause, A. Herbig, and C. Warinner, “The evolution and changing ecology of the African hominid oral microbiome,” Proceedings of the National Academy of Sciences, vol. 118, 5 2021.

[31] A. Fernandez-Guerra, G. Borrel, T. O. Delmont, B. Elberling, A. M. Eren, S. Gribaldo, A. Jochheim, R. A. Henriksen, K.-U. Hinrichs, T. S. Korneliussen, M. Krupovic, N. K. Larsen, R. Laso-Pérez, M. W. Pedersen, V. K. Pedersen, K. K. Sand, M. Sikora, M. Steinegger, I. Veseli, L. Wörmer, L. Zhao, M. Žure, K. Kjær, and E. Willerslev, “A 2-million-year-old microbial and viral communities from the Kap København Formation in North Greenland,” bioRxiv, 6 2023.

[32] D. Bozzi, S. Neuenschwander, D. I. Cruz Dávalos, B. Sousa da Mota, H. Schroeder, J. V. Moreno-Mayar, M. E. Allentoft, and A.-S. Malaspinas, “Towards predicting the geographical origin of ancient samples with metagenomic data,” Scientific Reports, vol. 14, Sept. 2024.

[33] M. A. Spyrou, K. I. Bos, A. Herbig, and J. Krause, “Ancient pathogen genomics as an emerging tool for infectious disease research,” Nature Reviews Genetics, vol. 20, p. 323–340, 4 2019.

[34] H. Al-Asadi, K. K. Dey, J. Novembre, and M. Stephens, “Inference and visualization of DNA damage patterns using a grade of membership model,” Bioinformatics, vol. 35, p. 1292–1298, 9 2018.

[35] L. Slimak, T. Vimala, A. Seguin-Orlando, L. Metz, C. Zanolli, R. Joannes-Boyau, M. Frouin, L. J. Arnold, M. Demuro, T. Deviese, D. Comeskey, M. Buckley, H. Camus, X. Muth, J. E. Lewis, H. Bocherens, P. Yvorra, C. Tenailleau, B. Duployer, H. Coqueugniot, O. Dutour, T. Higham, and M. Sikora, “Long genetic and social isolation in Neanderthals before their extinction,” Cell Genomics, vol. 4, p. 100593, Sept. 2024.

[36] A. Ginolhac, M. Rasmussen, M. T. P. Gilbert, E. Willerslev, and L. Orlando, “mapDamage: testing for damage patterns in ancient DNA sequences,” Bioinformatics, vol. 27, p. 2153–2155, June 2011.

[37] H. J ‘onsson, A. Ginolhac, M. Schubert, P. L. F. Johnson, and L. Orlando, “mapDamage2.0: fast approximate Bayesian estimates of ancient DNA damage parameters,” Bioinformatics, vol. 29, p. 1682–1684, Apr. 2013.

[38] P. Skoglund, B. H. Northoff, M. V. Shunkov, A. P. Derevianko, S. Pääbo, J. Krause, and M. Jakobsson, “Separating endogenous ancient DNA from modern day contamination in a Siberian Neandertal,” Proceedings of the National Academy of Sciences, vol. 111, p. 2229–2234, Jan. 2014.

[39] J. Neukamm, A. Peltzer, and K. Nieselt, “DamageProfiler: fast damage pattern calculation for ancient DNA,” Bioinformatics, vol. 37, p. 3652–3653, Apr. 2021.

[40] R. Everett and B. Cribdon, “MetaDamage tool: Examining post-mortem damage in sedaDNA on a metagenomic scale,” Frontiers in Ecology and Evolution, vol. 10, 1 2023.

[41] C. Michelsen, M. W. Pedersen, A. Fernandez-Guerra, L. Zhao, T. C. Petersen, and T. S. Korneliussen, “metaDMG – A Fast and Accurate Ancient DNA Damage Toolkit for Metagenomic Data,” bioRxiv, 12 2022.

[42] R. Hübler, F. M. Key, C. Warinner, K. I. Bos, J. Krause, and A. Herbig, “HOPS: automated detection and authentication of pathogen DNA in archaeological remains,” Genome Biology, vol. 20, Dec. 2019.

[43] Z. Pochon, N. Bergfeldt, E. Kirdök, M. Vicente, T. Naidoo, T. van der Valk, N. E. Altinişik, M. Krzewińska, L. Dalén, A. Götherström, C. Mirabello, P. Unneberg, and N. Oskolkov, “aMeta: an accurate and memory-efficient ancient metagenomic profiling workflow,” Genome Biology, vol. 24, Oct. 2023.

[44] A. Herbig, F. Maixner, K. I. Bos, A. Zink, J. Krause, and D. H. Huson, “MALT: Fast alignment and analysis of metagenomic DNA sequence data applied to the Tyrolean Iceman,” bioRxiv, Apr. 2016.

[45] Å. J. Vågene, A. Herbig, M. G. Campana, N. M. Robles García, C. Warinner, S. Sabin, M. A. Spyrou, A. Andrades Valtuenã, D. Huson, N. Tuross, K. I. Bos, and J. Krause, “Salmonella enterica genomes from victims of a major sixteenth-century epidemic in Mexico,” Nature Ecology & Evolution, vol. 2, p. 520–528, Jan. 2018.

[46] A. D. Williams, V. W. Leung, J. W. Tang, N. Hidekazu, N. Suzuki, A. C. Clarke, D. A. Pearce, and T. T.-Y. Lam, “Ancient environmental microbiomes and the cryosphere,” Trends in Microbiology, vol. 33, p. 233–249, Feb. 2025.

[47] A. Margaryan, D. J. Lawson, M. Sikora, F. Racimo, S. Rasmussen, I. Moltke, L. M. Cassidy, E. Jørsboe, A. Ingason, M. W. Pedersen, T. Korneliussen, H. Wilhelmson, M. M. Buś, P. de Barros Damgaard, R. Martiniano, G. Renaud, C. Bhérer, J. V. Moreno-Mayar, A. K. Fotakis, M. Allen, R. Allmäe, M. Molak, E. Cappellini, G. Scorrano, H. McColl, A. Buzhilova, A. Fox, A. Albrechtsen, B. Schütz, B. Skar, C. Arcini, C. Falys, C. H. Jonson, D. Blaszczyk, D. Pezhemsky, G. Turner-Walker, H. Gestsdóttir, I. Lundstrøm, I. Gustin, I. Mainland, I. Potekhina, I. M. Muntoni, J. Cheng, J. Stenderup, J. Ma, J. Gibson, J. Peets, J. Gustafsson, K. H. Iversen, L. Simpson, L. Strand, L. Loe, M. Sikora, M. Florek, M. Vretemark, M. Redknap, M. Bajka, T. Pushkina, M. Søvsø, N. Grigoreva, T. Christensen, O. Kastholm, O. Uldum, P. Favia, P. Holck, S. Sten, S. V. Arge, S. Ellingvåg, V. Moiseyev, W. Bogdanowicz, Y. Magnusson, L. Orlando, P. Pentz, M. D. Jessen, A. Pedersen, M. Collard, D. G. Bradley, M. L. Jørkov, J. Arneborg, N. Lynnerup, N. Price, M. T. P. Gilbert, M. E. Allentoft, J. Bill, S. M. Sindbæk, L. Hedeager, K. Kristiansen, R. Nielsen, T. Werge, and E. Willerslev, “Population genomics of the Viking world,” Nature, vol. 585, p. 390–396, Sept. 2020.

[48] E. I. Zavala, Z. Jacobs, B. Vernot, M. V. Shunkov, M. B. Kozlikin, A. P. Derevianko, E. Essel, C. de Fillipo, S. Nagel, J. Richter, F. Romagné, A. Schmidt, B. Li, K. O’Gorman, V. Slon, J. Kelso, S. Pääbo, R. G. Roberts, and M. Meyer, “Pleistocene sediment DNA reveals hominin and faunal turnovers at Denisova Cave,” Nature, vol. 595, p. 399–403, June 2021.

[49] A. Lien, L. P. Legori, L. Kraft, P. W. Sackett, and G. Renaud, “Benchmarking software tools for trimming adapters and merging next-generation sequencing data for ancient DNA,” Frontiers in Bioinformatics, vol. 3, Dec. 2023.

[50] F. Pedregosa, G. Varoquaux, A. Gramfort, V. Michel, B. Thirion, O. Grisel, M. Blondel, P. Prettenhofer, R. Weiss, V. Dubourg, J. Vanderplas, A. Passos, D. Cournapeau, M. Brucher, M. Perrot, and E. Duchesnay, “Scikit-learn: Machine Learning in Python,” Journal of Machine Learning Research, vol. 12, pp. 2825–2830, 2011.

[51] D. Li, C.-M. Liu, R. Luo, K. Sadakane, and T.-W. Lam, “MEGAHIT: an ultra-fast singlenode solution for large and complex metagenomics assembly via succinct de Bruijn graph,” Bioinformatics, vol. 31, p. 1674–1676, 1 2015.

[52] T. van der Valk, P. Pečnerová, D. Díez-del Molino, A. Bergström, J. Oppenheimer, S. Hartmann, G. Xenikoudakis, J. A. Thomas, M. Dehasque, E. Sağlican, F. R. Fidan, I. Barnes, S. Liu, M. Somel, P. D. Heintzman, P. Nikolskiy, B. Shapiro, P. Skoglund, M. Hofreiter, A. M. Lister, A. Götherström, and L. Dalén, “Million-year-old DNA sheds light on the genomic history of mammoths,” Nature, vol. 591, p. 265–269, Feb. 2021.

[53] L. Orlando, R. Allaby, P. Skoglund, C. Der Sarkissian, P. W. Stockhammer, M. C. Ávila Arcos, Q. Fu, J. Krause, E. Willerslev, A. C. Stone, and C. Warinner, “Ancient DNA analysis,” Nature Reviews Methods Primers, vol. 1, Feb. 2021.

