## Supplementary Material for "AdDeam: A Fast and Scalable Tool for Estimating and Clustering Reference-Level Damage Profiles"

**S3 Convergence Checking in `bam2prof`** 8

**S4 Runtime** 9

**S5 Additional information for Empirical Analysis: Mapping Ancient Metagenomic Fragments to Core Oral Microbiome Genera** 10

### S1 Generation of Simulated Data

To properly assess the clustering performance, we generated simulated sequencing reads where the ground truth—the true damage level of each sample—was known. Logically, all samples with the same simulated damage profile should be assigned to the same cluster. Moreover, the inferred damage profiles of the cluster should be similar to the original damage profile used for generating the samples of this cluster.

We simulated sequencing reads with `gargammel`[1] and aligned them to their respective reference genomes with `bwa aln`[2], resulting in 290 samples with no damage, 145 with mid damage, and 29 with high damage. The goal was to evaluate its ability to handle substitution frequencies that are nearly continuous, such as those ranging between 0.1 and 0.6. These situations are challenging because the patterns may not form visually distinct groups. Moreover, when analysing numerous profiles at the same time, manual cluster assignment quickly becomes impractical.

In empirical samples, damage levels vary between references, even among those that are expected to cluster together relative to other observable damage levels (e.g., one species may display 30% damage at the first position while another shows 27% or 35%). To simulate this empirical heterogeneity, we created three pools of 1,000 similar damage profiles corresponding to no damage, mid damage [3], and high damage [4]. Using 290 bacterial reference genomes, we simulated reads for each genome at all three damage levels by randomly sampling from the respective pools, resulting in a total of 870 datasets.

We simulated sequencing reads with `gargammel`[1] at an average coverage of 4X per genome. The resulting reads were then trimmed and merged using `leeHom`[5] with the `--ancientdna` parameter before being mapped to their respective reference genomes using `bwa aln`[2] with the parameters -n 0.01 -o 2 -l 1024 [6]. After alignment, we processed the BAM files with `samtools`[7] to sort them and add MD flags, which annotate mismatches relative to the reference.

We then processed these alignments using `AdDeam` in classic mode to generate damage profiles, resulting in 870 BAM files (290 per damage level). To better approximate empirical scenarios, we employed a non-uniform distribution of samples in our clustering analysis, using 290 samples with no damage, 145 samples with mid damage, and 29 samples with high damage. The damage profiles used for simulation and those estimated from the simulated data are shown in Figure S1, panel A.

The substitution frequencies in our simulations were designed to be distinct. However, the estimated damage profiles from aligned BAM files often differ from these true frequencies. This is due to biases in the alignment, introduced by the choice of reference and alignment parameters.

Fig. **S1** illustrates this situation. Panel A shows the simulated true damage profiles for each category. These represent substitution frequencies under ideal conditions - without reference bias

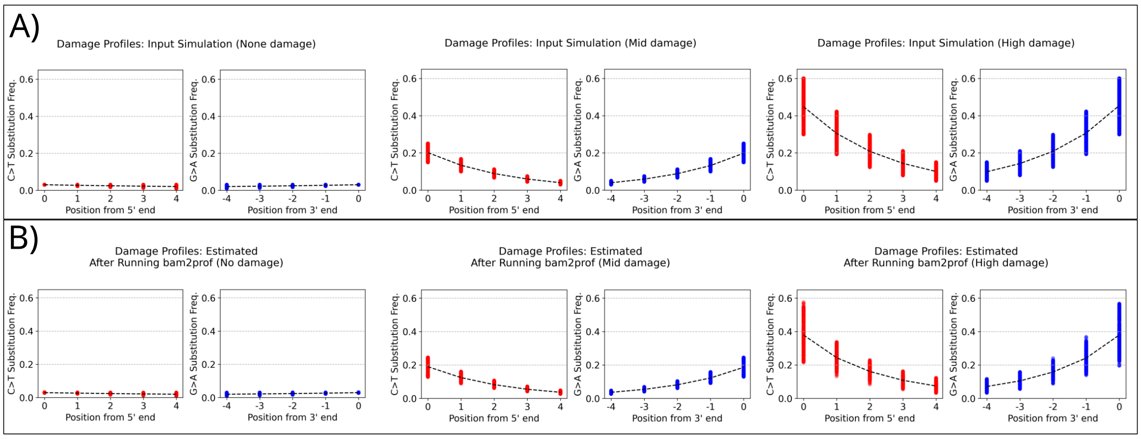

**Figure S1**: Simulated and estimated damage profiles. Panel A shows the simulated "true" substitution frequencies for each damage level. These represent ideal conditions without reference bias or misaligned reads. Panel B shows the estimated damage profiles after alignment and analysis.

or erroneously mapped reads. Of course, such conditions are impossible in real data because ambiguous mappings and global mapping parameters affect all reads equally, rather than being tailored to each read. Panel B shows the estimated damage profiles after alignment and analysis. Here, the separation between mid and high damage profiles is less clear, with significant overlap between the two categories.

This overlap highlights the value of clustering approaches like `AdDeam`. It groups similar profiles even when the distinctions are subtle or obscured by alignment biases. This becomes especially important when working with large datasets containing many damage profiles.

### S2 Comparison to similar tools

Benchmarking `AdDeam` against existing methods is challenging because other tools are designed with slightly different objectives in mind. The two most relevant tools are `PyDamage` [8], which assigns a probabilistic score to contigs reflecting their likelihood of ancient origin, and `aRchaic` [9], which clusters BAM files according to damage patterns. We outline the challenges of directly benchmarking `AdDeam` against these tools below.

With `AdDeam`, one can similarly identify contigs that are unlikely to be ancient by inspecting clusters with none or minimal damage signatures. Unlike `PyDamage`, which automatically filters out contigs that do not meet predefined model thresholds, `AdDeam` presents all cluster assignments to the user and leaves the final interpretation to them. Overall, `PyDamage` assigns each contig a model-inferred score for a more quantitative, albeit less transparent, classification. In contrast, `AdDeam` clusters contigs into a user-specified number of $k$ clusters, providing an intuitive visualisation rather than a strict, model-based binary cut-off. Therefore, direct comparison of the two tools is of limited applicability, as they employ fundamentally different approaches.

aRchaic reports a "grade of membership" for each sample, defined as the fraction of reads assigned to each damage profile [9]. In contrast, AdDeam first computes a single aggregated damage profile per BAM (or per reference), then applies a Gaussian mixture model to these profiles to obtain probabilistic cluster assignments. Crucially, these probabilities reflect the similarity of the entire damage profile to each cluster centroid, not the proportion of reads exhibiting a particular pattern. For instance, a sample with both ancient and modern contamination might receive membership scores of approximately 80% "ancient" and 20% "modern" in aRchaic, whereas AdDeam would derive one intermediate profile and assign it to the cluster whose centroid best matches that mixed-damage signature. Therefore, the GMM-based probabilities in AdDeam and the membership grades in aRchaic measure fundamentally different aspects of damage, making direct comparison difficult.

Nevertheless, to benchmark AdDeam, we compared its performance against PyDamage [8] and aRchaic [9]. In addition, we evaluated the impact of alternative clustering algorithms on our results.

##### S2.1 Comparison to aRchaic

Using the original aRchaic datasets from Al-Asadi *et al.* [9], which include No-UDG samples (Skoglund *et al.* [10], Gamba *et al.* [11]), UDG samples Lipson *et al.* [12], Lazaridis *et al.* [13] and modern samples from the 1000 Genomes project [14]. Since the aRchaic paper does not reveal the exact sample identifiers, we collected subsamples of similar sizes. All sample identifiers are listed in the data repository under https://github.com/LouisPwr/AdDeamAnalysis.

We ran AdDeam using the default -lib paired flag. Although the Gamba *et al.* [11] libraries were single-end, their blunt-end repair produced the classic C→T and G→A signatures. With $k = 3$ (compare to Figure 2 in [9]), we recovered identical clusters: one containing all modern samples, one with non-UDG libraries, and one with UDG-treated samples (Figure **S2**).

The main difference lies in the cluster probability assignments. In aRchaic, grades of membership reveal that ancient samples retain a residual modern signal with UDG-treated samples, more so than non-UDG, whereas AdDeam generates near-binary assignments, with each sample showing ≈100% probability for its assigned cluster.

While the cluster assignments were identical, aRchaic's grade-of-membership approach more directly highlights contamination levels. In this dataset, AdDeam's cluster probabilities appear almost binary: Each sample is assigned with approximately 100% confidence, likely because samples within each cluster are highly homogeneous and markedly distinct from those in other clusters.

A less binary behaviour emerged in the analysis of the *de novo* assembled contigs (Main Text,

Figure 4), where many contigs receive split probabilities well below 100% for their primary cluster. Note that the individual damage patterns are also much more evenly distributed, as can be seen from the numerous grey traces in Figure 4.

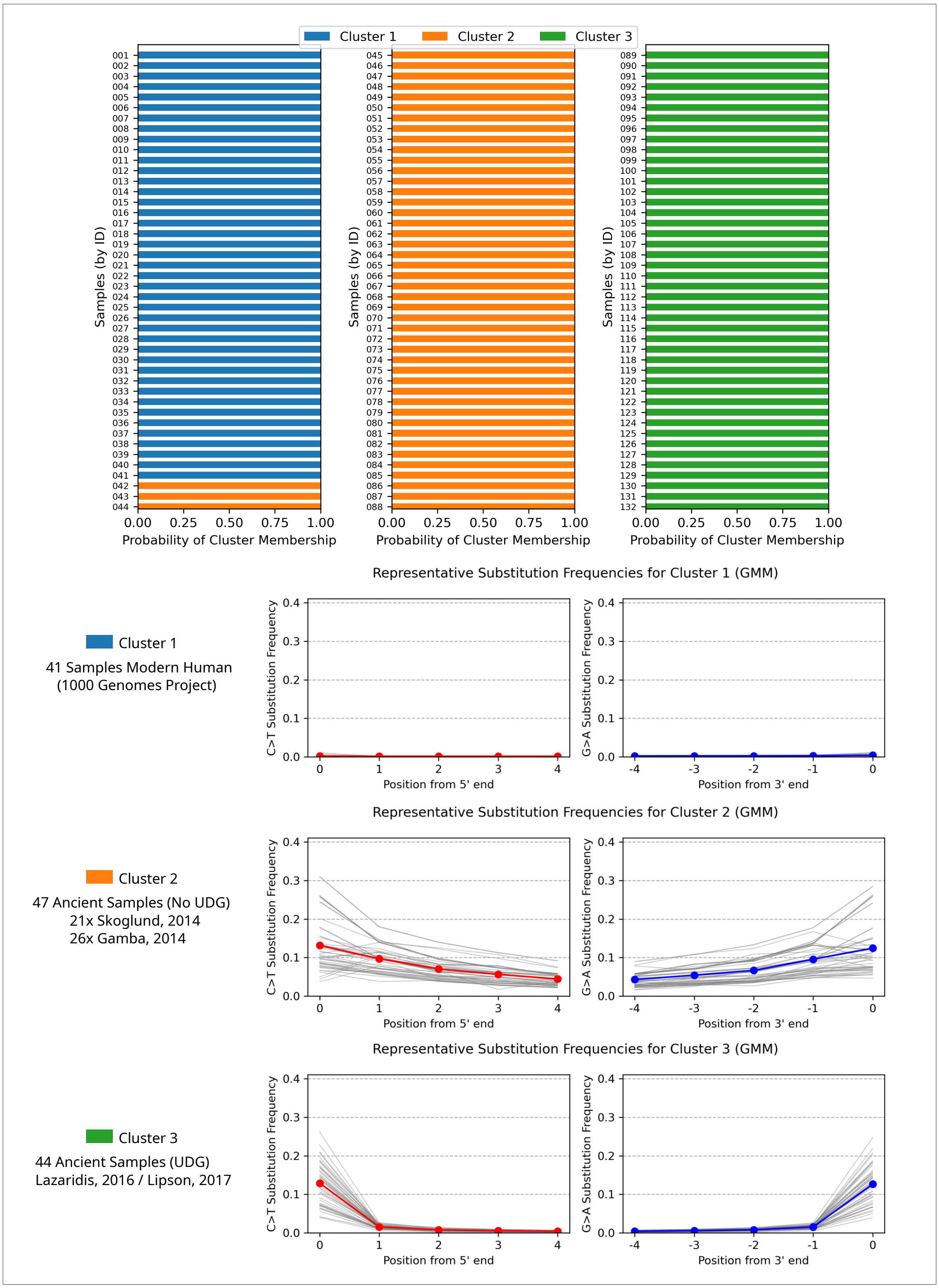

**Figure S2**: Top: Probability of each sample belonging to each of the three clusters inferred by `AdDeam`. Bottom: Damage-profile plots for each cluster, showing the cluster representative (bold line) and individual sample profiles (fine grey lines). Sample identifiers for each cluster are annotated at the left of the corresponding profile.

#### S2.2 Comparison to PyDamage

We compared `AdDeam` and `PyDamage` by analysing the same BAM file containing alignments to the *de novo* assembled contigs used in Main Text, Section "4.3 Empirical Analysis: Clustering Damage Profiles from Assembled Contigs." `PyDamage` was run with default filters ($q$-value>0.05 and predicted_accuracy<0.5) to partition contigs into "likely ancient" and "unlikely ancient" sets. In parallel, `AdDeam` produced four sets of contigs: those whose damage profiles did not converge (and thus were not clustered) and the contigs assigned to Clusters 1, 2, and 3 (see Main Text, Figure 4 for representative profiles). Cluster 1 contains contigs with minimal substitution frequencies, Clusters 2 and 3 progressively higher frequencies.

Figure **S3** presents an Upset plot of all intersections among these five sets. For the "unlikely ancient" group, `PyDamage Not Ancient` and `AdDeam Cluster 1` share 114 contigs, and `PyDamage Not Ancient` and `AdDeam Not Converged` share 99. No contigs in `AdDeam` Clusters 2 or 3 appear in the `PyDamage Not Ancient` set, in agreement with the elevated damage rates in these clusters.

By contrast, the "likely ancient" contigs show more divergence. `PyDamage Ancient` overlaps 697 contigs in `AdDeam Cluster 1`, 517 in Cluster 2, 87 in Cluster 3, and 256 in the Not Converged set. This pattern suggests that `AdDeam` is more conservative when applied for labelling contigs as ancient. In particular, many Cluster 1 contigs derive 98.8% of their reads from the modern sample and exhibit extremely low substitution frequencies (Main Text Figure 4), yet they are nonetheless called ancient by `PyDamage`, implying that its significance thresholds may be too permissive in this context.

Overall, both tools concordantly identified a core set of non-ancient contigs, but differed in sensitivity. Consequently, `PyDamage` and `AdDeam` can be used complementarily: `PyDamage` offers a quantitative per-contig framework, while `AdDeam`'s cluster-representative profiles can guide the selection or adjustment of significance thresholds.

#### S2.3 Alternative Clustering Approaches

Clustering results inevitably depend on the chosen algorithm. In developing `AdDeam`, we evaluated several probabilistic and distance-based methods—including non-negative matrix factorization (NMF), soft/fuzzy $k$-means, agglomerative hierarchical clustering, dynamic time warping (DTW)–based clustering, and Gaussian mixture models (GMM; see Main Text, Section 3.2). We tested each on simulated datasets with varying proportions of no, mid-level, and high damage. Only the spherical-covariance GMM and DTW-based clustering perfectly recovered the known clusters.

Despite the appeal of modelling position-to-position correlations, a spherical-covariance GMM

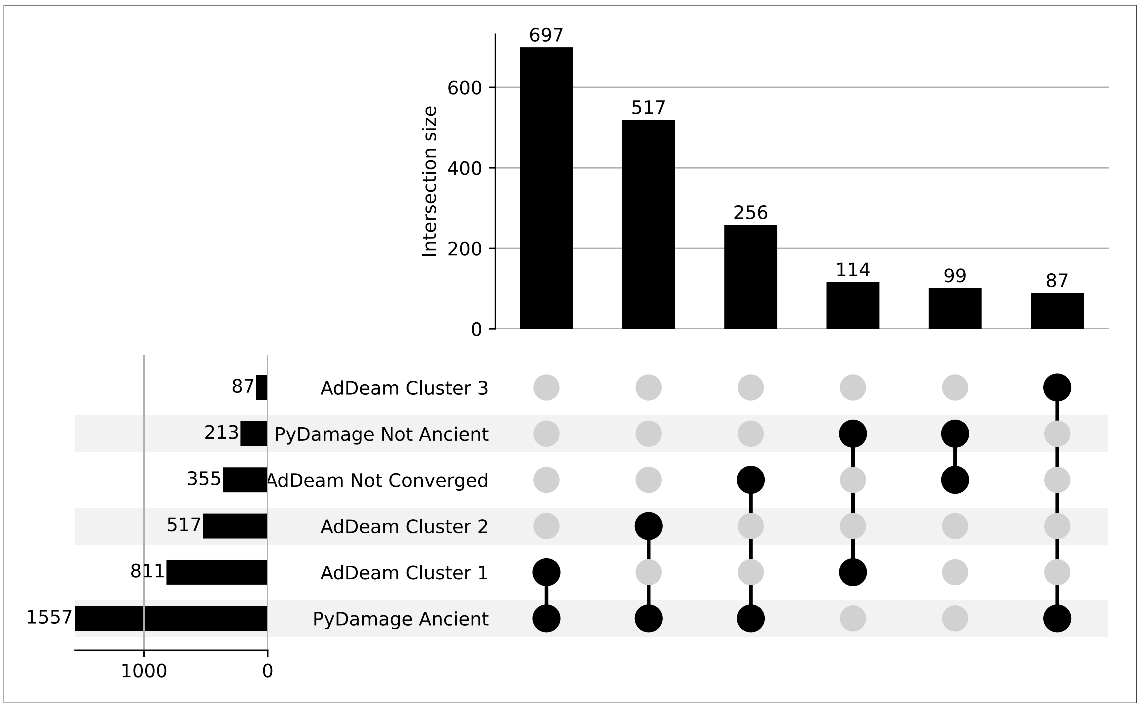

**Figure S3**: Upset plot comparing contig classifications by `PyDamage` and `AdDeam`. **Left bars:** Total number of contigs in each set - "PyDamage Ancient," "PyDamage Not Ancient," "AdDeam Not Converged," and Clusters 1–3. **Matrix:** Connected dots indicate which sets participate in each intersection. **Top bars:** Size of each intersection, i.e., the number of contigs shared among the specified sets. This layout highlights the strong concordance on non-ancient contigs (e.g., overlap between `PyDamage Not Ancient` and `AdDeam Cluster 1`/Not Converged) and the more variable agreement for ancient contigs.

proved most robust. Although spherical covariances treat each read position as independent (i.e., they reduce to Euclidean distances in feature space), using a full covariance matrix led to overfitting and more incorrect clusters on both simulated and empirical data.

DTW offers an alternative by non-linearly "warping" the read-position axis to align length-$p$ damage trajectories (e.g. matching shifted damage peaks). In a DTW $k$-medoids framework, each cluster centre is an actual profile (medoid), which proved computationally efficient in our tests. For $k = 3$, DTW recovered the same cluster assignments as the GMM on the `aRchaic` test set (Figure S2). However, for $k = 2$, DTW clustered five UDG-treated samples together with the modern samples, likely because DTW prioritizes local slope similarity over the overall damage-profile shape.

Soft-DTW $k$-means, by contrast, requires iterative barycentre-averaging and proved ≈70x slower than the spherical GMM on our 1,770 contig profiles (Main Text, Figure 4), making it impractical for large datasets. Thus, although DTW methods can in principle leverage sequential dependencies, neither variant outperformed the spherical-covariance GMM in both speed and clustering accuracy for `AdDeam`.

Overall, DTW-based clustering did not outperform the spherical-covariance GMM in our evaluation, but it showed promise for damage-profile clustering and could be explored further in future work.

#### S3 Convergence Checking in `bam2prof`

`bam2prof`, the subprogram in `AdDeam` that computes the damage profiles, implements an iterative convergence check to both (1) accelerate damage profile computation and (2) identify problematic "zig-zag" patterns that can skew clustering results. After every $n$ processed alignments, `bam2prof` computes the absolute change in substitution frequencies (e.g., C→T) over the last $n$ reads. If this change falls below a user-defined threshold $d$, the program terminates early, avoiding the need to process all fragments. Figure **S4**, panel **(A)**, illustrates this behavior for the sample `Gok2cop_CGATGT_L007` (Skoglund *et al.* 2014; ENA project PRJEB6090), where the C→T differences every 500 fragments converge after roughly 10,000 reads. Both the check interval $n$ (`-numAligned`) and threshold $d$ (`-precision`) can be configured by the user.

In "meta" mode, the same convergence criterion flags profiles that never stabilize, e.g. due to low coverage or mis-mappings, by writing them to a separate directory with the suffix *_notConverged*. Our experience showed that using $n = 500$ and $d = 0.01$ effectively filters out non-converging "zig-zag" profiles. Figure **S4**, panel **B**, shows an example from sample OAK003 (Fellows-Yates *et al.* 2021; ENA project PRJEB34569), where C→T differences remain large up to 1,000 reads and converge quickly towards $\sim$ 10,000 processed fragments.

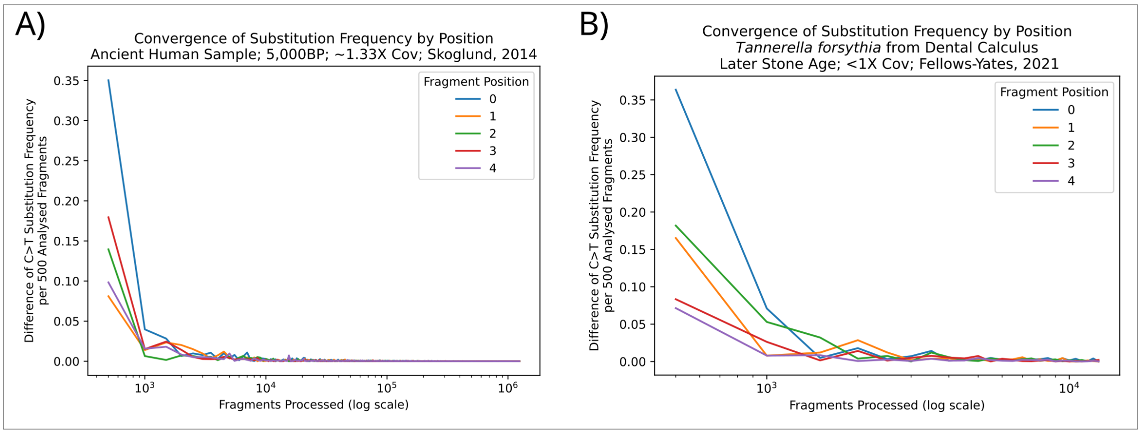

**Figure S4**: Convergence of C→T substitution–frequency differences as the number of processed fragments increases, evaluated every 500 fragments. **(A)** Line plot of the absolute change in C→T frequency over successive 500-read windows for an ancient human sample (`Gok2cop_CGATGT_L007`, Skoglund et al. 2014). Each colored trace corresponds to a different read-position (0–4) and shows how frequency changes diminish as more fragments are processed. **(B)** Equivalent plot for a metagenomic sample (OAK003, Fellows-Yates et al. 2021), highlighting persistent fluctuations at low fragment counts.

### S4 Runtime

We measured the runtime of each `AdDeam` processing step during the analysis of the *de novo* assembled contigs from empirical datasets (Section "Empirical Analysis: Clustering Damage Profiles from Assembled Contigs", Main). For a BAM file containing 1,770 references (contigs) and a total of 48,312,572 mapped fragments, the `bam2prof` module generated all 1,770 damage profiles in under 34 seconds. The clustering module completed clustering for $k = 2, 3$, and 4 in 1 minutes and 30 seconds, while running in quiet mode (i.e., with reduced visual output, beneficial for larger datasets) reduced the execution time to 45 seconds. The metrics were measured on a Intel(R) Xeon(R) Platinum 8462Y+ server with a "Processor Base Frequency" of 2.10 GHz and a "Max Turbo Frequency" of 4.10 GHz. The maximum memory usage was less than 400 Mb. Both damage profile generation and clustering utilized a single thread only. The `addeam-bam2prof.py` wrapper offers a `-hpc` option that redirects commands to a file, which can then be executed on HPC clusters that often use queueing systems.

### S5 Additional information for Empirical Analysis: Mapping Ancient Metagenomic Fragments to Core Oral Microbiome Genera

Table S1 summarizes the samples clustered in Main Text Section 4.2, Figure 3. For each sample, it reports the assigned cluster, sample code, host species (dental calculus source), archaeological period, and UDG treatment status.

| **Cluster Assigned** | **Sample codes** | **Sample Source** | **Period** | **UDG Treated** |
|---|---|---|---|---|
| 1 | DJA (1x) | Gorilla | 20th century | yes |
| 1 | EBO (1x) | Gorilla | 20th century | yes |
| 1 | OME (1x) | Howler Monkey | 20th-21st century | no |
| 1 | OME (3x) | Howler Monkey | 20th-21st century | yes |
| 1 | OAK (2x) | Modern human | Later Stone Age | yes |
| 1 | TAF (1x) | Modern human | Later Stone Age | yes |
| 1 | ECO (2x) | Modern human | Mesolithic | yes |
| 1 | FUM (1x) | Neanderthal | Middle Palaeolithic | yes |
| 1 | PES (1x) | Neanderthal | Middle Palaeolithic | yes |
| 1 | EMN (1x) | Modern human | Upper Palaeolithic | yes |
| 2 | DJA (5x) | Chimpanzee | 20th century | no |
| 2 | EBO (6x) | Chimpanzee | 20th century | no |
| 2 | OME (3x) | Howler Monkey | 20th-21st century | no |
| 3 | OAK (3x) | Modern human | Later Stone Age | no |
| 3 | TAF (4x) | Modern human | Later Stone Age | no |
| 3 | FUM (3x) | Neanderthal | Middle Palaeolithic | no |
| 3 | PES (1x) | Neanderthal | Middle Palaeolithic | no |
| 3 | EMN (1x) | Modern human | Upper Palaeolithic | no |
| 3 | ECO (1x) | Modern human | Mesolithic | no |
| 4 | OAK (1x) | Modern human | Later Stone Age | no |
| 4 | TAF (2x) | Modern human | Later Stone Age | no |
| 4 | ECO (5x) | Modern human | Mesolithic | no |

Table S1: Overview of sample information and cluster assignments. The table includes sample codes, sources, time periods, and whether the samples were UDG-treated.

### References

[1] G. Renaud, K. Hanghøj, E. Willerslev, and L. Orlando, "gargammel: a sequence simulator for ancient DNA," Bioinformatics, vol. 33, p. 577–579, 11 2016.

[2] H. Li and R. Durbin, "Fast and accurate short read alignment with Burrows–Wheeler transform," Bioinformatics, vol. 25, p. 1754–1760, May 2009.

[3] I. Olalde, M. E. Allentoft, F. Sánchez-Quinto, G. Santpere, C. W. K. Chiang, M. DeGiorgio, J. Prado-Martinez, J. A. Rodríguez, S. Rasmussen, J. Quilez, O. Ramírez, U. M. Marigorta, M. Fernández-Callejo, M. E. Prada, J. M. V. Encinas, R. Nielsen, M. G. Netea, J. Novembre, R. A. Sturm, P. Sabeti, T. Marquès-Bonet, A. Navarro, E. Willerslev, and C. Lalueza-Fox, "Derived immune and ancestral pigmentation alleles in a 7, 000-year-old Mesolithic European," Nature, vol. 507, p. 225–228, Jan. 2014.

[4] T. Günther, H. Malmström, E. M. Svensson, A. Omrak, F. Sánchez-Quinto, G. M. Kılınç, M. Krzewińska, G. Eriksson, M. Fraser, H. Edlund, A. R. Munters, A. Coutinho, L. G. Simões, M. Vicente, A. Sjölander, B. Jansen Sellevold, R. Jørgensen, P. Claes, M. D. Shriver, C. Valdiosera, M. G. Netea, J. Apel, K. Lidén, B. Skar, J. Storå, A. Götherström, and M. Jakobsson, "Population genomics of Mesolithic Scandinavia: Investigating early postglacial migration routes and high-latitude adaptation," PLOS Biology, vol. 16, p. e2003703, Jan. 2018.

[5] G. Renaud, U. Stenzel, and J. Kelso, "leeHom: adaptor trimming and merging for Illumina sequencing reads," Nucleic Acids Research, vol. 42, p. e141–e141, 8 2014.

[6] A. Oliva, R. Tobler, A. Cooper, B. Llamas, and Y. Souilmi, "Systematic benchmark of ancient DNA read mapping," Briefings in Bioinformatics, vol. 22, Apr. 2021.

[7] H. Li, B. Handsaker, A. Wysoker, T. Fennell, J. Ruan, N. Homer, G. Marth, G. Abecasis, and R. Durbin, "The Sequence Alignment/Map format and SAMtools," Bioinformatics, vol. 25, p. 2078–2079, 6 2009.

[8] M. Borry, A. Hübner, A. B. Rohrlach, and C. Warinner, "PyDamage: automated ancient damage identification and estimation for contigs in ancient DNA de novo assembly," PeerJ, vol. 9, p. e11845, July 2021.

[9] H. Al-Asadi, K. K. Dey, J. Novembre, and M. Stephens, "Inference and visualization of DNA damage patterns using a grade of membership model," Bioinformatics, vol. 35, p. 1292–1298, 9 2018.

[10] P. Skoglund, H. Malmström, A. Omrak, M. Raghavan, C. Valdiosera, T. Günther, P. Hall, K. Tambets, J. Parik, K.-G. Sjögren, J. Apel, E. Willerslev, J. Storå, A. Götherström, and

M. Jakobsson, "Genomic Diversity and Admixture Differs for Stone-Age Scandinavian Foragers and Farmers," Science, vol. 344, p. 747–750, May 2014.

[11] C. Gamba, E. R. Jones, M. D. Teasdale, R. L. McLaughlin, G. Gonzalez-Fortes, V. Mattiangeli, L. Domboróczki, I. Kővári, I. Pap, A. Anders, A. Whittle, J. Dani, P. Raczky, T. F. G. Higham, M. Hofreiter, D. G. Bradley, and R. Pinhasi, "Genome flux and stasis in a five millennium transect of European prehistory," Nature Communications, vol. 5, Oct. 2014.

[12] M. Lipson, A. Szécsényi-Nagy, S. Mallick, A. Pósa, B. Stégmár, V. Keerl, N. Rohland, K. Stewardson, M. Ferry, M. Michel, J. Oppenheimer, N. Broomandkhoshbacht, E. Harney, S. Nordenfelt, B. Llamas, B. Gusztáv Mende, K. Köhler, K. Oross, M. Bondár, T. Marton, A. Osztás, J. Jakucs, T. Paluch, F. Horváth, P. Csengeri, J. Koós, K. Sebők, A. Anders, P. Raczky, J. Regenye, J. P. Barna, S. Fábián, G. Serlegi, Z. Toldi, E. Gyöngyvér Nagy, J. Dani, E. Molnár, G. Pálfi, L. Márk, B. Melegh, Z. Bánfai, L. Domboróczki, J. Fernández-Eraso, J. Antonio Mujika-Alustiza, C. Alonso Fernández, J. Jiménez Echevarría, R. Bollongino, J. Orschiedt, K. Schierhold, H. Meller, A. Cooper, J. Burger, E. Bánffy, K. W. Alt, C. Lalueza-Fox, W. Haak, and D. Reich, "Parallel palaeogenomic transects reveal complex genetic history of early European farmers," Nature, vol. 551, p. 368–372, Nov. 2017.

[13] I. Lazaridis, D. Nadel, G. Rollefson, D. C. Merrett, N. Rohland, S. Mallick, D. Fernandes, M. Novak, B. Gamarra, K. Sirak, S. Connell, K. Stewardson, E. Harney, Q. Fu, G. Gonzalez-Fortes, E. R. Jones, S. A. Roodenberg, G. Lengyel, F. Bocquentin, B. Gasparian, J. M. Monge, M. Gregg, V. Eshed, A.-S. Mizrahi, C. Meiklejohn, F. Gerritsen, L. Bejenaru, M. Blüher, A. Campbell, G. Cavalleri, D. Comas, P. Froguel, E. Gilbert, S. M. Kerr, P. Kovacs, J. Krause, D. McGettigan, M. Merrigan, D. A. Merriwether, S. O'Reilly, M. B. Richards, O. Semino, M. Shamoon-Pour, G. Stefanescu, M. Stumvoll, A. Tönjes, A. Torroni, J. F. Wilson, L. Yengo, N. A. Hovhannisyan, N. Patterson, R. Pinhasi, and D. Reich, "Genomic insights into the origin of farming in the ancient Near East," Nature, vol. 536, p. 419–424, July 2016.

[14] The 1000 Genomes Project Consortium, "A global reference for human genetic variation," Nature, vol. 526, p. 68–74, Sept. 2015.
